# Genome-wide screen for genes required for smooth lipopolysaccharide production in Escherichia coli K-12

**DOI:** 10.64898/2026.07.26.740846

**Authors:** Jilong Qin, Elizabeth Ngoc Hoa Tran, Vincenzo Leo, Yaoqin Hong, Alistair J. Standish, Renato Morona

**Affiliations:** Institute for Molecular Bioscience, The University of Queensland, Brisbane, QLD; Australia School of Biological Sciences, Research Centre for Infectious Diseases, University of Adelaide, Adelaide 5005, Australia; Australian Institute of Tropical Health and Medicine, James Cook University, Townsville, Queensland, Australia; Division of Biomedicine and Molecular Biology, College of Medicine and Dentistry, James Cook University, Townsville, Queensland, Australia; College of Medicine and Public Health, Flinders University, Bedford Park 5042, South Australia, Australia

**Keywords:** KEIO collection, O-antigen, LPS, Colicin E2, *E*. *coli* K-12

## Abstract

Lipopolysaccharide (LPS) is a major component of the outer membrane of Gram-negative bacteria, contributing to membrane integrity and environmental interactions. Genome-wide studies defining bacterial gene functions have been extensively performed in the model strain *Escherichia coli* K-12 which lacks O-antigen (OAg), and therefore does not produce smooth LPS (S-LPS). Consequently, the genetic requirements for S-LPS production in this model system remain incompletely defined. Here, a functional *wbbL* gene was introduced into the *E. coli* K-12 KEIO single-gene deletion mutant library to restore OAg synthesis, enabling genome-wide analysis of S-LPS production by screening with colicin E2 (ColE2) and validated with LPS silver staining. This identified 319 mutants with increased sensitivity to ColE2 in the presence of OAg, suggesting broader envelope-associated effects during screening. In addition, 27 mutants showed defects in S-LPS production, corresponding to genes involved in OAg biosynthesis, LPS core and sugar precursor synthesis, OAg ligation and regulation, and enterobacterial common antigen biosynthesis. A further 18 mutants initially appeared defective in S-LPS production but could not be validated upon reconstruction, and whole-genome sequencing revealed secondary mutations responsible for the observed phenotypes. This study provides a validated genetic framework for S-LPS production in *E. coli* K-12 and highlights the importance of rigorous validation in genome-wide screening approaches.

## Introduction

The outer membrane is a defining feature of Gram-negative bacteria and serves as an essential permeability barrier that protects cells against environmental stresses, antimicrobial compounds, and host immune defences [1]. A major structural component of the outer membrane is lipopolysaccharide (LPS), which contributes to membrane integrity, outer membrane asymmetry, and interactions between bacteria and their surrounding environment [2]. The LPS molecule is composed of three structurally distinct regions: lipid A, the core oligosaccharide, and the O antigen (OAg). Depending on the presence or absence of the OAg, LPS exists as either smooth LPS (S-LPS) or rough LPS (R-LPS). The OAg forms the outermost layer of the bacterial surface and plays important roles in protection against complement-mediated killing [3], resistance to bacteriophages [4], and adaptation to diverse environmental conditions [5]. In pathogenic bacteria, the OAg is also a major virulence determinant [6] and an important target for host immune recognition [7].

The biosynthesis of the OAg is a highly coordinated process that involves numerous enzymes responsible for the synthesis of nucleotide sugar precursors, assembly of repeating oligosaccharide units on the undecaprenyl phosphate lipid carrier, translocation across the inner membrane, polymerisation of repeating units, and ligation of the completed polysaccharide to the lipid A-core molecule [8]. In *Escherichia coli*, the Wzx/Wzy-dependent pathway is the predominant mechanism for OAg biosynthesis, requiring the coordinated activities of glycosyltransferases, the Wzx flippase, the Wzy polymerase, the Wzz chain-length regulator, and the WaaL ligase [9]. While the core enzymatic steps of OAg assembly have been extensively characterised, efficient production of S-LPS also depends on proper synthesis of the LPS core [10], maintenance of lipid A-core assembly [11], adequate availability of activated sugar precursors [12], and coordination with other envelope glycoconjugate biosynthetic pathways, such as enterobacterial common antigen (ECA) [13]. Perturbations in these processes can indirectly impair either OAg polymerisation or ligation, leading to defective S-LPS production despite otherwise intact OAg biosynthetic machinery.

Although *E. coli* K-12 has long served as one of the principal model organisms for bacterial genetics and physiology, it does not produce an OAg because of genetic inactivation of the *wbbL* gene within its O16 OAg biosynthetic gene cluster [14]. The *wbbL* gene encodes a rhamnosyltransferase required for synthesis of the OAg repeating unit, and complementation with a functional copy of *wbbL* restores the ability of K-12 strains to produce S-LPS. This genetic background provides a powerful experimental platform for investigating the cellular pathways required for OAg biosynthesis while retaining the extensive genetic resources available for *E. coli* K-12.

The KEIO collection [15], comprising nearly all non-essential single-gene deletion mutants of *E. coli* K-12, has become an invaluable resource for systematic functional genomic studies. Genome-wide screening of this collection enables systematic assessment of the contribution of individual non-essential genes to complex cellular processes and has proven valuable for defining the genetic basis of diverse aspects of bacterial physiology. However, despite considerable advances in understanding the enzymatic pathway of OAg biosynthesis, relatively few studies have systematically assessed the contribution of individual non-essential genes to S-LPS production at a genome-wide scale. Consequently, the complete set of non-essential genes required for efficient S-LPS expression in *E. coli* K-12 remains to be experimentally defined.

In this study, we employed a genome-wide genetic approach to identify genes required for S-LPS production by introducing a functional *wbbL* gene into the *E. coli* K-12 KEIO mutant collection and screening with colicin E2 (ColE2) for mutants displaying defects in S-LPS production. This systematic analysis provides a comprehensive evaluation of the non-essential genes required for efficient S-LPS production in *E. coli* K-12 and defines the genetic determinants necessary for OAg synthesis, ligation and expression on the bacterial cell surface.

## Results

### *An engineered mobilisable* wbbL *clone restored O16-OAg production of BW25113*

The *E. coli* K-12 strain lacks OAg production due to its genetic inactivation in the gene encoding committed-step glycosyltransferase WbbL [14, 16]. In order to restore the production of OAg of the entire KEIO library [17] (3909 mutants based on *E. coli* K-12 strain BW25113) through high-throughput conjugation, we cloned *wbbL* into a mobilisable IncP-type vector pSUP203 [18] for constitutive expression, designated pWbbL. As expected, complementation of wild-type (WT) BW25113 by pWbbL restored O16 OAg production (Fig 1A). The production of OAg impacts the accessibility of ColE2, a DNA exonuclease [19] to its cellular receptor BtuB in the outer membrane, thereby abolishing the uptake of ColE2 [20] and conferring resistance [21]. As a result, the restoration of O16 OAg in BW25113 by complementing *wbbL* allowed it to grow on LB agar medium supplemented with purified ColE2 (LBA-ColE2) (Fig 1A). These results confirmed that constitutive expression of *wbbL* in BW25113 restores O16 OAg production which confers resistance to ColE2 for subsequent screening of OAg production defects.

**Fig 1.**
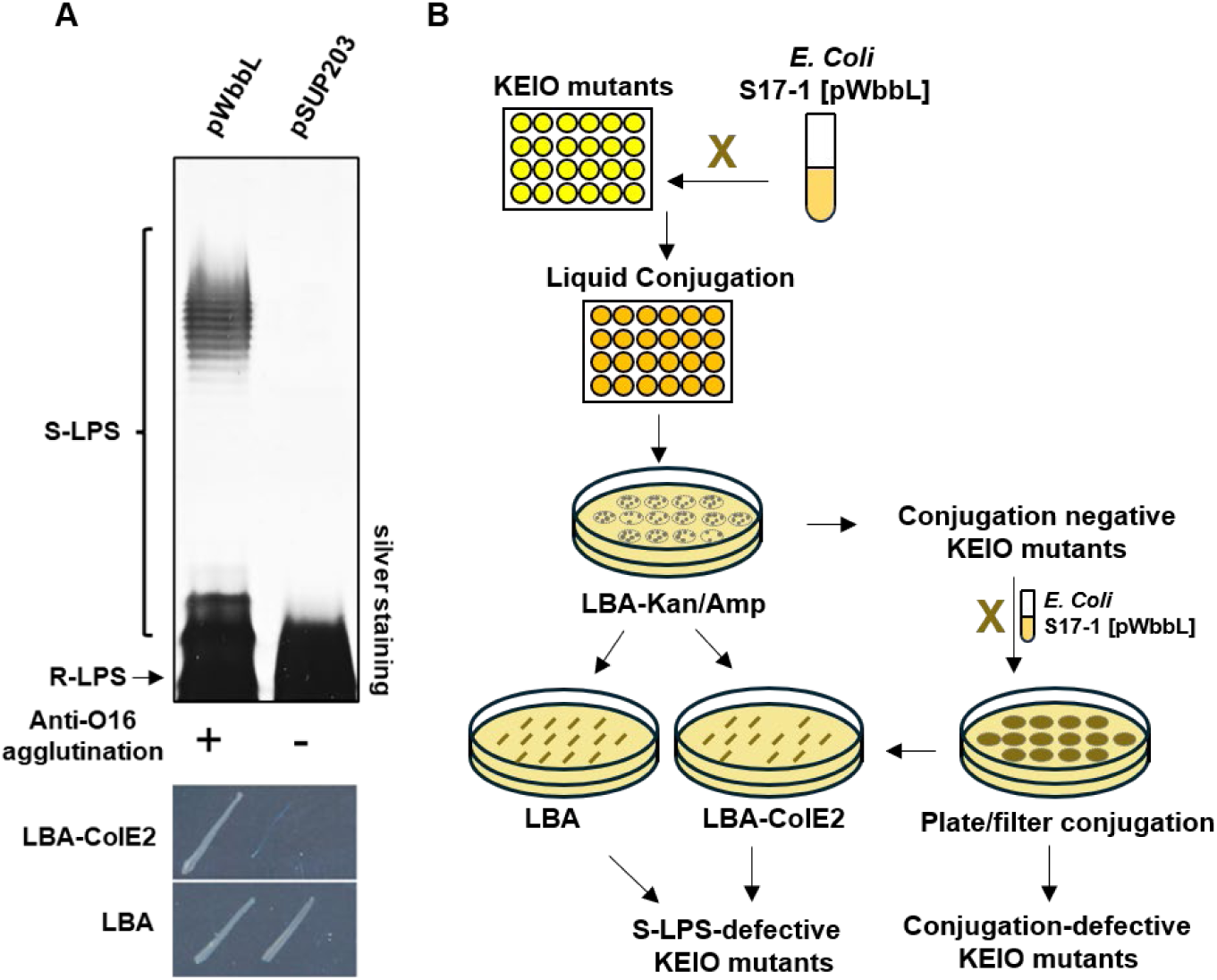
Genome-wide screening of the *E. coli* KEIO knockout library via ColE2 susceptibility following restoration of OAg biosynthesis. A) Complementation of *wbbL* in BW2113 restores O16 OAg S-LPS production and confer resistance to ColE2. Proteinase K-treated whole cell lysate was analysed via SDS-PAGE and silver staining. The presence of O16 OAg was confirmed by agglutination with anti-O16 antiserum. Bacterial colonies were streaked onto LB agar plates supplemented with ColE2 for sensitivity validation. B) Schematic overview of complementation of whole KEIO library using high throughput conjugation, following by ColE2 selection, and phenotypic characterisation workflow.

### *High-throughput conjugation to complement* wbbL *of entire KEIO library revealed conjugation-defective mutants*

To restore the OAg biosynthesis pathway in all KEIO mutants, we next introduced pWbbL into a mobilising strain (S17-1) engineered with chromosomally integrated conjugative IncP-type RP4 [18] as a donor strain (S17-1 [pWbbL]) (Fig 1B). Among the 3,909 KEIO mutants, three mutants, *ybeY* and *stfQ*, which did not result in growth, and *yhbG*, which was previously confirmed with partial duplication [22], were excluded from the high-throughput conjugation. The remaining KEIO mutants were then mated with *E. coli* S17-1[pWbbL] in liquid media to allow conjugation, and the resulting transconjugants were selected on LB agar plates supplemented with appropriate antibiotics (LBA-Kan/Amp) (Fig 1B). Of the liquid conjugation, 3,821 mutants successfully yielded transconjugants, while 85 mutants failed to receive pWbbL through conjugation in liquid broth (Table S1). This group includes: 1) the OAg flippase *wzxB* (*rfbX*), which has been shown previously to be lethal when OAg production is restored in *E. coli* K-12 due to the sequestration of UndP in UndPP-OAg intermediates [23]. This result confirms that our high-throughput conjugation is robust enough to identify the lethality caused by single gene deletions in the OAg biosynthesis pathway when OAg production is restored; 2) genes responsible for the synthesis of lipopolysaccharide core oligosaccharide, *gmhA* (*lpcA*), *waaQ* (*rfaQ*) and *waaG* (*rfaG*), which were reported previously to impact the conjugation efficiency [24]; 3) genes responsible for plasmid DNA replication including *dnaT*; 4) genes that belong to the *tol-pal* operon including *tolR, tolB* and *pal,* which affects beta-lactam resistance [25], ultimately affecting transconjugants selection on ampicillin-containing media, and 5) gene disruptions that result in poor growth of the recipient strain, which would give a conjugation-deficient phenotype, such as *rimM*, *rnt* and *lapB* (*yciM*) [26].

Since conjugation efficiency is higher on solid surfaces for RP4 encoding rigid pili [27], we next performed conjugation on solid media (LBA) surfaces for the remaining 84 mutants excluding the *wzxB* mutant. However, 6 single gene deletion mutants only yielded colonies when the conjugation mixture was concentrated (5×) and plated (Table S1), this included *rnt, yfbK, yfdH, rfe (wecA), ycjU* and *ygeN,* suggesting that these gene functions may impact conjugation efficiency. Despite showing sensitivity to ColE2 (Table S1), the mutants with low plate conjugation efficiency were excluded from further investigation to avoid ColE2 sensitivity effects resulting from potential selection for suppressor mutations during conjugation. Nevertheless, all the remaining gene mutants yielded transconjugants and were included in the subsequent S-LPS production screening. In contrast to *gmhA*, *waaG* and *lapB,* which all showed increased sensitivity to ColE2, *waaQ* remained resistant to ColE2 upon WbbL complementation.

### ColE2 screening revealed non-essential genes required for S-LPS production

Overall, we identified 319 ColE2 sensitive mutants (CSM) upon *wbbL* complementation (Table S2). To determine whether the decrease in ColE2 sensitivity was due to a defect in lipopolysaccharide (LPS), the LPS profiles of all 319 CSMs complemented with *wbbL* were then analysed by SDS-PAGE and silver staining (Table S2). CSMs showing an impaired S-LPS profile (compared to the parent BW25113 complemented with *wbbL*) were observed for 45 gene knock-outs (Table S2). Eighteen mutants with no prior known association with LPS biosynthesis showed defective S-LPS profiles after been complemented by pWbbL (Table S2). However, the defect of these mutants in S-LPS production could not be validated as they showed S-LPS profiles when these knock-out mutants were remade in the *E. coli* K-12 background complemented with pWbbL (Table S3). We then selected 8 of these KEIO mutants for whole genome sequencing (WGS), and confirmed secondary mutations including *wzyB*, *waaB*, *wbbJ*, *gmhB*, *glf* and *waaC* in gene products affecting S-LPS biosynthesis (Table S3). Interestingly, we found secondary mutations in *aceE* in both *ycbU* and *ybjJ* KEIO mutants, yet the role of *aceE* in S-LPS is unclear. Collectively, these data suggest that the defective phenotype in S-LPS production in these 18 KEIO mutants are due to secondary mutations found in S-LPS biogenesis. The remaining 27 mutants are known to impact polysaccharide biogenesis (Table 1). These are known genes products responsible for i) the biogenesis of OAg repeating units and their assembly (Fig. 2) including *rmlD*, *rmlC*, *wbbK, wbbI*, *wbbJ* and *glf* which showed R-LPS; as well as *rmlA*, *wzzB* and *yfdI*, which showed reduced S-LPS, unregulated and shifted S-LPS modality, respectively; ii) the LPS core oligosaccharide and assembly (Fig 3) including *waaP, waaR, rfaD, waaF, waaO, galU, galE, hldE, waaH, gmhB, waaC, waaB, gmhA* and *waaG*, which all showed truncations in LPS core oligosaccharide, termed deep-rough LPS (DR-LPS), except for *waaH*, which only showed to have impaired S-LPS production quantity; iii) the LPS biogenesis regulation, *rfaH* and *lapB*; iv) the OAg ligase *waaL*; and v) ECA biogenesis *wecD* (*rffC*). Taken together, our data successfully screened out non-essential gene products affecting S-LPS production in *E. coli* K-12.

**Fig 2.**
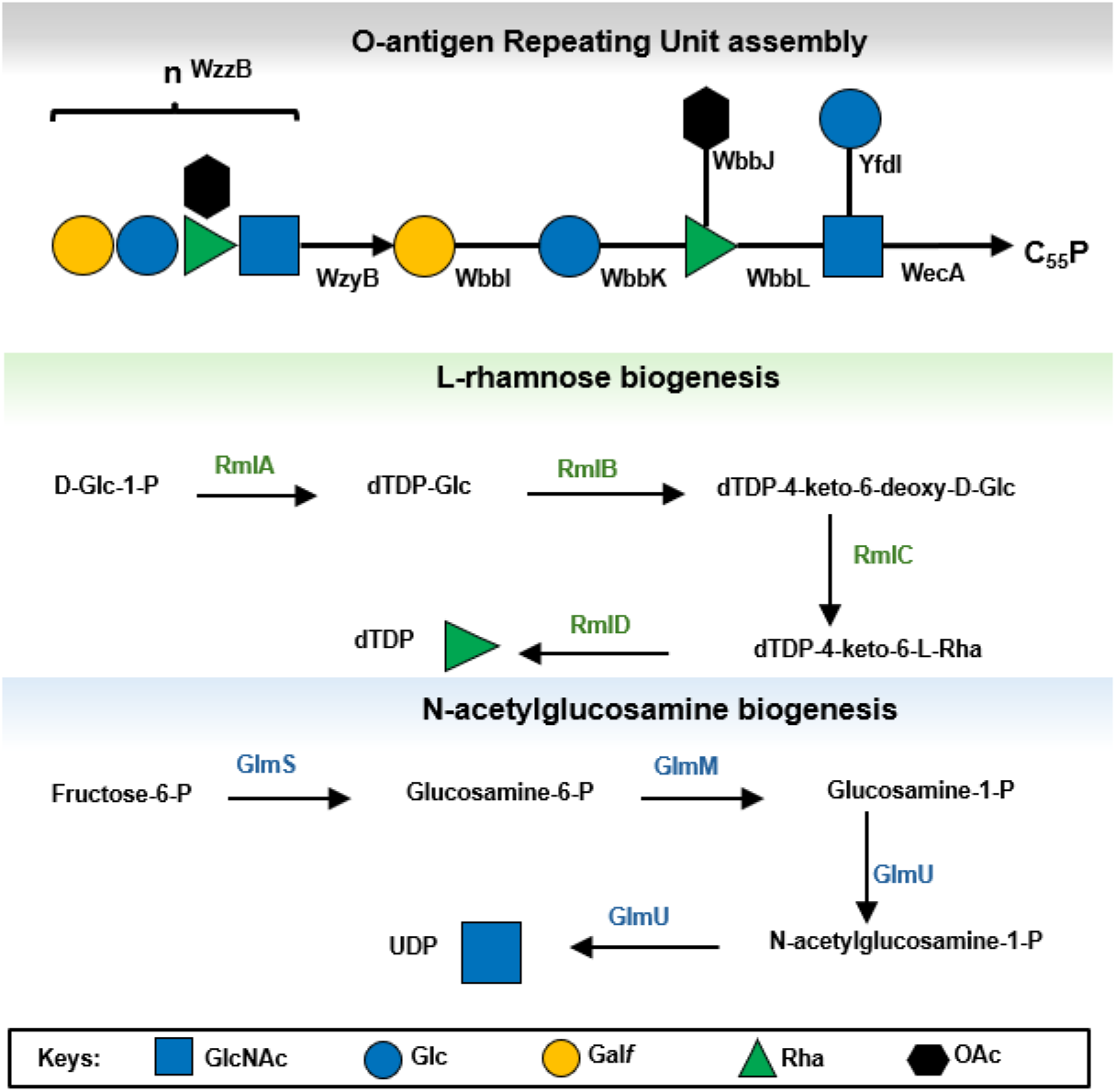
Schematic representation of OAg repeating unit biosynthesis and assembly. Sugar precursors are sequentially assembled onto undecaprenyl phosphate by a series of glycosyltransferases, generating the OAg repeating unit. Individual glycosyltransferases involved in core oligosaccharide assembly are indicated. The lower panel summarise the biosynthetic pathways for key precursor sugars.

**Fig 3.**
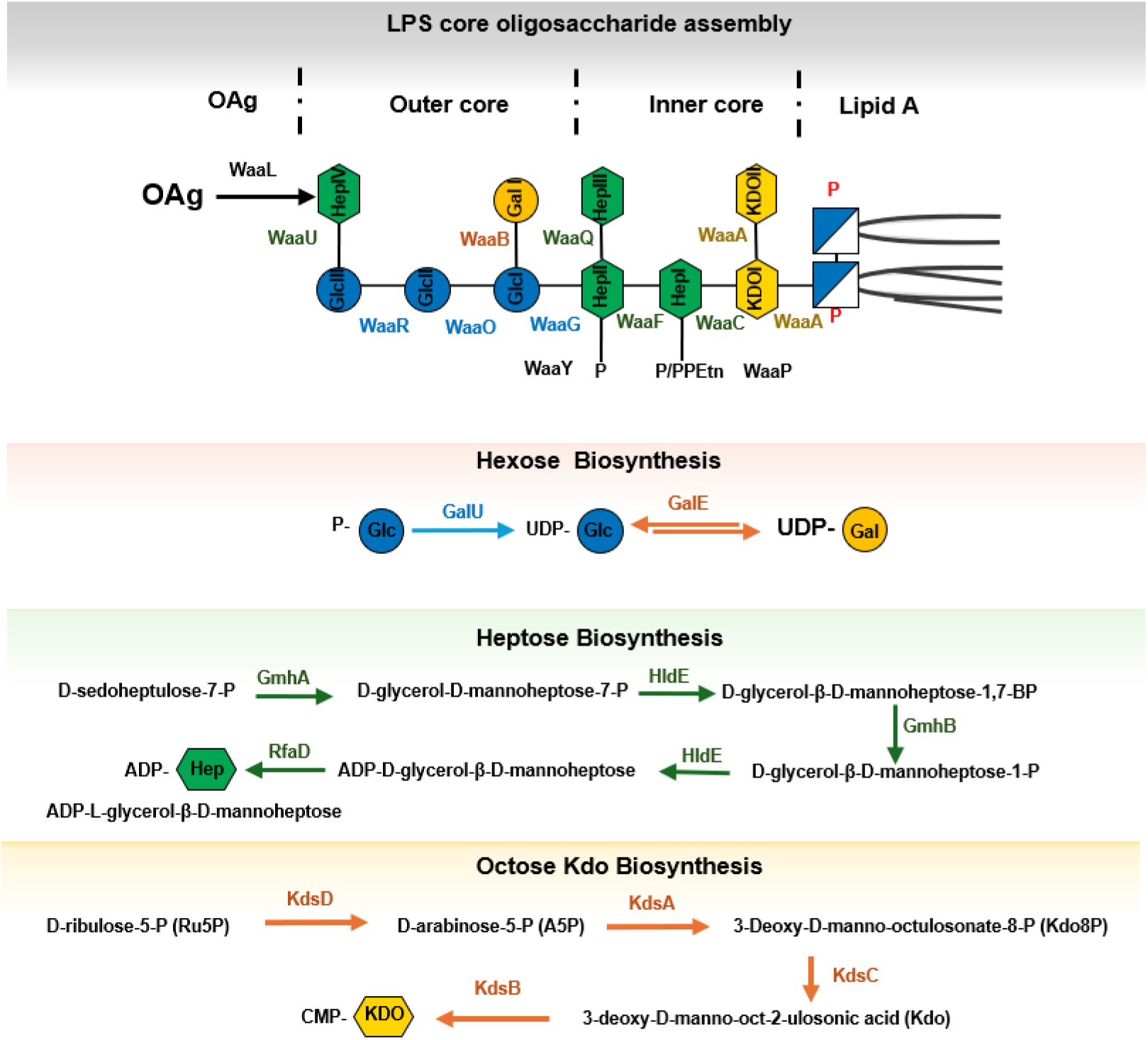
Schematic representation of LPS core oligosaccharide biosynthesis and assembly. The upper panel depicts the stepwise addition of sugar residues to form the inner and outer core regions, followed by ligation of the OAg to generate S-LPS. Individual glycosyltransferases involved in core oligosaccharide assembly are indicated. The lower panels summarise the biosynthetic pathways for key LPS core precursor sugars, including hexose, heptose, and 3-deoxy-D-manno-octulosonic acid (Kdo) synthesis.

**Table 1.**
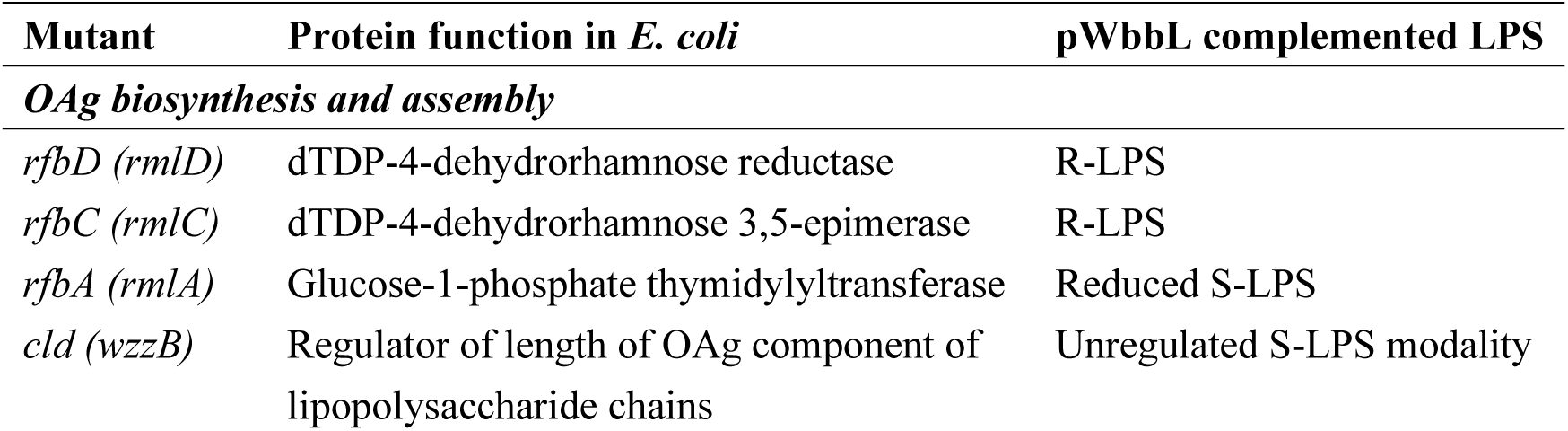
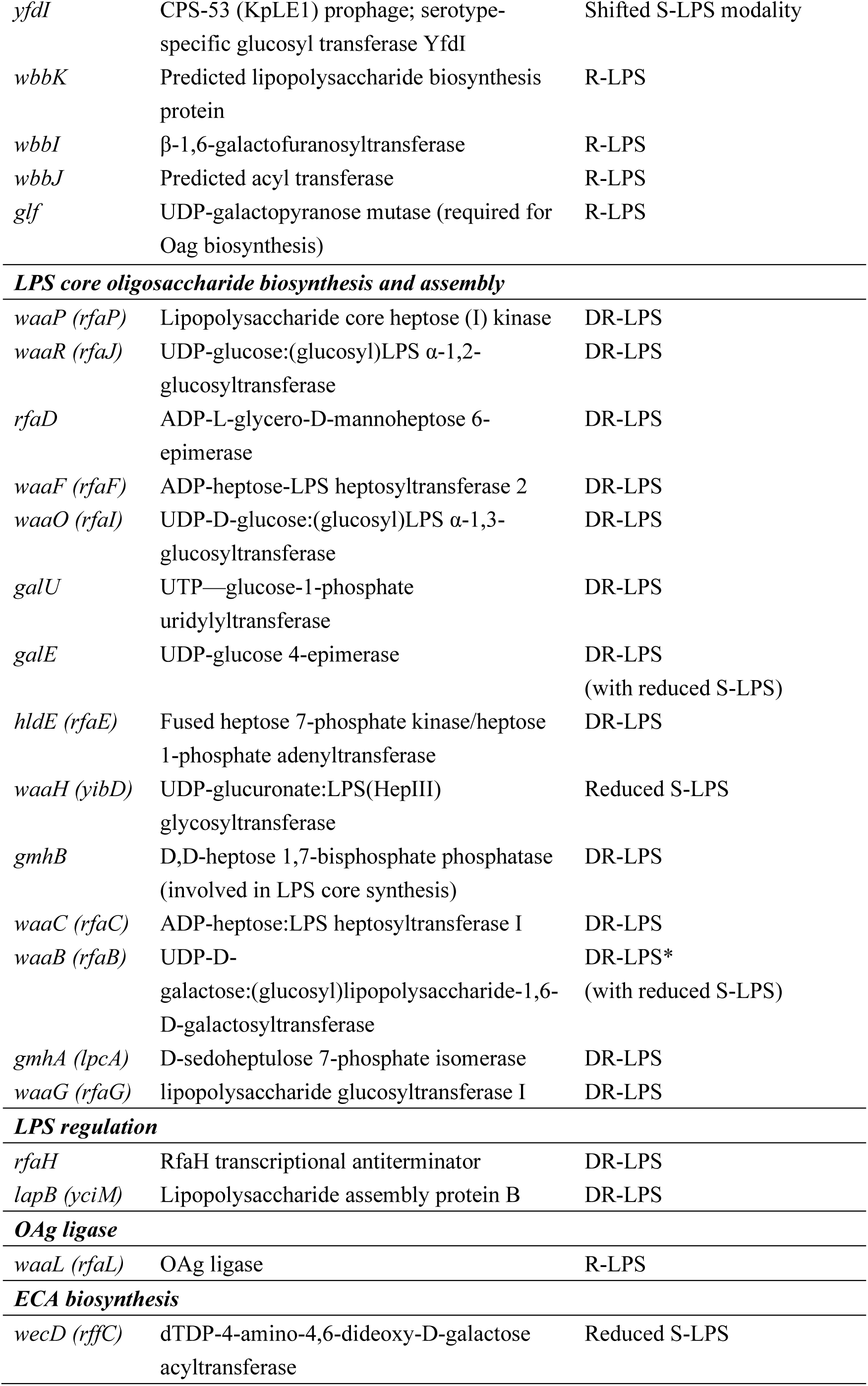
Non-essential genes responsible for S-LPS production.

## Discussion

This study provides a genome-wide assessment of non-essential genes required for S-LPS production in *Escherichia coli* K-12 complemented with *wbbL*. The majority of identified genes belong to well-established pathways of OAg biosynthesis, LPS core assembly, and nucleotide sugar precursor production, confirming that S-LPS biogenesis primarily depends on the coordinated activity of canonical envelope biosynthetic machineries. These findings provide a validated genetic framework for S-LPS production in a model organism that is otherwise widely used for genome-wide functional studies.

A key limitation of the KEIO-based approach is that it is restricted to non-essential genes and does not account for genetic redundancy. Multiple pathways involved in envelope biogenesis contain paralogous or compensatory enzymes, such that loss of a single gene may not produce an observable S-LPS phenotype. For example, redundancy in nucleotide sugar biosynthesis pathways, including KDO metabolism mediated by *kdsC* and *kdsD* [28], may buffer against single-gene perturbations, masking potential contributors to S-LPS assembly. Similarly, genes involved in envelope maintenance or regulation may only manifest phenotypes under specific environmental conditions not captured in standard laboratory growth. Indeed, *rfbB* (*rmlB*), a gene involved in dTDP-rhamnose biosynthesis for OAg assembly, displayed a S-LPS phenotype in our screen. However, previous studies in *Salmonella* have shown that disruption of *rfbB* alone does not completely abolish OAg production, unless *rffG* (*rmlB* involved in ECA-related glycan precursor metabolism) is also disrupted [29]. This suggests that nucleotide sugar metabolism and envelope glycoconjugate pathways may exhibit compensatory or interconnected effects that influence S-LPS production. In contrast, a gene associated with ECA biosynthesis, *wecD* (*rffC*), was identified as affecting S-LPS production, whereas most other ECA pathway genes, including *wzxE* and *wecF*, were not recovered. Although ECA and OAg biosynthesis share membrane-linked intermediates, including the undecaprenyl phosphate carrier, the absence of broader ECA pathway hits suggests that ECA does not generally contribute to OAg assembly or stabilisation under conditions of restored S-LPS production. Instead, the effect of *wecD* may reflect a specific influence on envelope glycoconjugate homeostasis, precursor allocation, or carrier lipid availability rather than functional redundancy between the ECA and OAg systems.

Metabolic background also strongly influences S-LPS production. The identification of *galE* highlights the dependence of OAg synthesis on nucleotide sugar availability. In *galE*-deficient conditions, depletion of UDP-galactose can lead to a mixed population of truncated LPS species, including deep-rough phenotypes, depending on growth composition and exogenous galactose sugar availability [30, 31]. This underscores the sensitivity of S-LPS assembly to metabolic flux and nutrient conditions.

The screen also highlighted the importance of rigorous validation in genome-wide envelope studies. A subset of initial hits, particularly those associated with conjugation-related functions, likely reflected indirect effects arising from the experimental delivery system used for *wbbL* introduction [24]. In contrast, several apparent S-LPS defects were not directly attributable to the targeted gene deletions. For example, the S-LPS-defective phenotype observed in an *oppF* mutant was resolved upon reconstruction and was found to be caused by a secondary mutation in *wzyB,* consistent with its essential role in OAg polymerization and S-LPS production. Notably, no *wzyB* mutant was recovered as a true positive in the initial S-LPS screen. This observation suggests that the *wzyB* mutant in our KEIO library may be incorrect. Consistent with this, whole-genome sequencing of selected mutants confirmed that 18 initially identified S-LPS-defective strains carried secondary mutations responsible for the observed phenotypes. Together, these findings emphasise the necessity of genetic complementation and whole-genome validation when interpreting envelope-related phenotypes in high-throughput screening approaches.

The identification of *lapB* as a potential contributor to S-LPS production may be explained by its indirect effects on LPS biogenesis. First, LapB has been shown to regulate LpxC stability and influence LPS synthesis, and perturbation of this pathway can affect Lpt-dependent LPS transport processes [32]. Since LpxC participates in the biosynthesis of lipid A portion of the R-LPS, which serves as the acceptor substrate for OAg ligation to generate S-LPS, changes in LapB activity may indirectly alter S-LPS levels. Second, *lapB* deletion causes a severe growth defect at 37°C [32], the temperature used for the screen, and this impaired envelope homeostasis or reduced cellular fitness may further affect the efficiency of S-LPS production. In contrast, the deletion of LPS-related genes such as *rfaQ* showed normal S-LPS profiles, which is consistent with previous studies [33], suggests that not all LPS-associated genes are essential for S-LPS production under laboratory conditions.

Our screen highlights differences in assay sensitivity. LPS silver staining primarily reflects bulk abundance of LPS species and may not capture subtle structural or spatial defects in OAg distribution. In contrast, ColE2 sensitivity proved to be a more sensitive indicator of S-LPS perturbations, likely reflecting changes in surface accessibility rather than total LPS quantity [21]. These observations suggest that spatial organisation and dynamic assembly of LPS may be as important as total production levels in determining envelope functionality, which warrant a future study. Together, these results define the non-essential genetic requirements for S-LPS production in *E. coli* K-12 and highlight both the robustness and limitations of genome-wide knockout approaches.

## Materials and Methods

### Bacterial strains and maintenance

The strains used in this study were *Escherichia coli* K-12 BW25113, S17-1 [18] and the KEIO collection mutants [15]. The *E. coli* K-12 KEIO collection was obtained from the National BioResource Project (NIG, Japan) which includes *E. coli* (total 3,909 mutants) on 50 rectangular agar plates containing 96 bacterial dots/plate. Once received, the entire collection was replicated in 1.2 ml deep well trays (LLG # 9407509) containing LB (pH 7.5) supplemented with 15 % (v/v) glycerol and kanamycin using a 96-pin microplate replicator (Boekel Scientific #140500). Trays were then sealed (Eppendorf mats #0030127552) and incubated at 37°C for 16 h with aeration, followed by storage at -80°C. Strains were routinely cultured in Lysogeny broth (LB) at 37°C with aeration. Where required, media were supplemented with 50 µg/ml kanamycin (Kan), 10 µg/ml tetracycline (Tet) or 100 µg/ml ampicillin (Amp).

### Construction of plasmids

The conjugative plasmid pSUP203-WbbL was made by PCR amplifying the *wbbL* encoding region from pMF19 [34] with oligos (GGCGCGGAATTCACTGAAGAACATTGAAATGGTAT and CGCCCCGAATTCTGCAGCTCGCGGCCTGGAATG), and ligating the EcoRI-digested fragment into pSUP203 [18]. A pSUP205-WbbL (a tetracycline resistant plasmid derivative) was made by sub-cloning the EcoRI-digested *wbbL* fragment from pSUP203-WbbL into pSUP205 [18].

### Serum agglutination assay

*E. coli* antisera O16 was purchased from SSI Diagnostica (#SSI45755). Agglutination of bacteria by antisera was assessed by emulsifying fresh bacterial growth from LB agar (4-6 colonies) in 40 μl PBS and mixing 1:1 with antisera. Positive agglutination was determined by eye and light microscopy if it occurred within 15 s.

### LPS PAGE and silver staining

Bacterial cells (10^9^ CFU) were lysed in 50 μl of SDS sample buffer, followed by heating at 100 °C for 10 minutes. The samples were then treated with 50 μg/ml proteinase K (PK, NEB) for 18 h at 60 °C. Three microliter of each sample was loaded onto 15% SDS PAGE gels, with LPS being silver stained as previously described [35].

### ColE2 purification and susceptibility assay

Purification of ColE2 and spot sensitivity assays were performed as previously described [16, 21]. Briefly, C-terminal his-tagged ColE2 protein with its immunity protein were expressed from BL21(DE3) carrying pET41b-ColE2 [36] and purified by immobilised metal affinity chromatography (IMAC) using Profinity Ni-charged resin (Bio-Rad) according to manufacturer’s protocol. For ColE2 susceptibility assays, strains grown to mid-exponential phase (OD ∼0.5) were spread onto LB agar, and 5 μl of purified ColE2 diluted in 2-fold series in PBS were spotted onto the plate. Plates were incubated overnight at 37°C for 18 h and the sensitivity level was determined by the minimum ColE2 concentration that showed clear bacterial growth inhibition.

### High-throughput bacterial conjugation and LPS defective screen

To perform the high throughput bacterial conjugation, 96 well trays containing 100 µl LB were systematically inoculated with the KEIO collection mutants, trays were parafilmed and incubated for 16 h at 37°C with aeration. SL17-1[pWbbL] (donor strain) was inoculated into 10 ml LB and incubated for 16 h at 37°C with aeration. SL17-1[pWbbL] was washed twice and diluted 1:2 in LB. Conjugations were then performed by mixing 10 µl of diluted SL17-1[pWbbL] culture with the 16 h KEIO cultures grown in the 96 well tray and left at 37°C for 5-6 h. Conjugation mixes (3 µl) were then dotted onto LB agar supplemented with Kan and Amp, following by incubation at 37°C for 18 h, and the growth from dots patched onto 15 ml LB agar (with Kan and Amp) supplemented with and without 2 µg/ml ColE2. Following a further overnight incubation at 37°C, the growth of conjugates in the presence and absence of ColE2 was noted.

### Standard bacterial plate conjugations

For mutants that failed to yield transconjugants, donor and recipient strains were grown for 16 h at 37℃ with aeration, washed twice in LB and mixed in a 1:10 ratio. Mating mixtures (5 ml) were then pelleted by centrifugation, resuspended in 50 µl LB, and pipetted onto sterile cellulose acetate membrane filters (0.45 μm, type HA, Millipore) placed centrally on a pre-warmed (37°C) LB plate. After 4 h incubation at 37°C, bacteria were collected off the filters by transferring the membranes to 5 ml LB broth (or 1 ml where indicated) and vortexing. The cell suspension (3 µl) was then dotted onto selective agar, and incubated at 37°C for 16 h. Growth from conjugate dots were then patched and incubated as above to select for ColE2 sensitive mutants (CSM).

### Mutagenesis via allelic exchange

*E. coli* and *S. flexneri* gene inactivation mutants were generated by Lambda Red mutagenesis as described previously [37]. Briefly, antibiotic resistance cassette was generated via PCR with oligos flanked with sequences homologous to targeted genes and was then used for allelic exchange gene replacement. Successful gene replacement as confirmed with targeted PCR.

### Whole genome sequencing analysis

Bacterial whole genome sequencing analysis were performed as described previously [16]. Briefly, genomic DNA *E. coli K-12* mutants were prepared using a Qiagen QIAamp DNA blood mini kit according to the manufacturer’s protocol. Samples were prepared for DNBseq DNA library construction (BGI) followed by DNBSEQ PE150 sequencing (BGI). The processed reads for each strain were then mapped onto the NCBI BW25113 reference genome (Accession number NZ_CP009273) using Geneious.

## Author contributions

JQ, RM contributed to the project conceptualisation; JQ, RM and ET contributed to experimental design; JQ, ET, VL, and AS contributed to data collection and analysis; JQ, ET and RM contributed to data interpretation; RM contributed to the core materials and the funding. YH contributed to the generation of experimental materials. JQ and ET drafted the manuscript, and all authors edited the manuscript.

## Data availability

All data generated or analysed during this study were included in this article and supplementary files.

## References

1. Nikaido H. Molecular basis of bacterial outer membrane permeability revisited. Microbiology and molecular biology reviews : MMBR. 2003;67(4):593–656. Epub 2003/12/11. doi: 10.1128/MMBR.67.4.593-656.2003. PubMed PMID: 14665678; PubMed Central PMCID: PMCPMC309051.

2. Raetz CR, Whitfield C. Lipopolysaccharide endotoxins. Annual review of biochemistry. 2002;71:635–700. doi: 10.1146/annurev.biochem.71.110601.135414. PubMed PMID: 12045108; PubMed Central PMCID: PMC2569852.

3. Murray GL, Attridge SR, Morona R. Altering the length of the lipopolysaccharide O antigen has an impact on the interaction of Salmonella enterica serovar Typhimurium with macrophages and complement. J Bacteriol. 2006;188(7):2735–9. doi: 10.1128/JB.188.7.2735-2739.2006. PubMed PMID: 16547065; PubMed Central PMCID: PMCPMC1428429.

4. Lindberg AA, Sarvas M, Makela PH. Bacteriophage attachment to the somatic antigen of salmonella: effect of o-specific structures in leaky R mutants and s, t1 hybrids. Infect Immun. 1970;1(1):88–97. doi: 10.1128/iai.1.1.88-97.1970. PubMed PMID: 16557701; PubMed Central PMCID: PMCPMC415860.

5. Reeves PR. Variation in O-antigens, niche-specific selection and bacterial populations. FEMS Microbiol Lett. 1992;100(1-3):509–16. doi: 10.1111/j.1574-6968.1992.tb14085.x. PubMed PMID: 1282486.

6. Pluschke G, Mercer A, Kusecek B, Pohl A, Achtman M. Induction of bacteremia in newborn rats by Escherichia coli K1 is correlated with only certain O (lipopolysaccharide) antigen types. Infect Immun. 1983;39(2):599–608. doi: 10.1128/iai.39.2.599-608.1983. PubMed PMID: 6187683; PubMed Central PMCID: PMCPMC347994.

7. Reeves P. Role of O-antigen variation in the immune response. Trends Microbiol. 1995;3(10):381–6. doi: 10.1016/s0966-842x(00)88983-0. PubMed PMID: 8564356.

8. Samuel G, Reeves P. Biosynthesis of O-antigens: genes and pathways involved in nucleotide sugar precursor synthesis and O-antigen assembly. Carbohydrate research. 2003;338(23):2503–19. Epub 2003/12/13. doi: 10.1016/j.carres.2003.07.009. PubMed PMID: 14670712.

9. Hong Y, Hu D, Verderosa AD, Qin J, Totsika M, Reeves PR. Repeat-Unit Elongations To Produce Bacterial Complex Long Polysaccharide Chains, an O-Antigen Perspective. EcoSal Plus. 2023:eesp00202022. Epub 2023/01/10. doi: 10.1128/ecosalplus.esp-0020-2022. PubMed PMID: 36622162.

10. Raetz CR, Reynolds CM, Trent MS, Bishop RE. Lipid A modification systems in gram-negative bacteria. Annual review of biochemistry. 2007;76:295–329. doi: 10.1146/annurev.biochem.76.010307.145803. PubMed PMID: 17362200; PubMed Central PMCID: PMCPMC2569861.

11. Mettlach JA, Cian MB, Chakraborty M, Dalebroux ZD. Signaling through the Salmonella PbgA-LapB regulatory complex activates LpxC proteolysis and limits lipopolysaccharide biogenesis during stationary-phase growth. J Bacteriol. 2024;206(4):e0030823. Epub 20240327. doi: 10.1128/jb.00308-23. PubMed PMID: 38534107; PubMed Central PMCID: PMCPMC11025326.

12. Weiner IM, Higuchi T, Osborn MJ, Horecker BL. Biosynthesis of O-antigen in Salmonella typhimurium. Annals of the New York Academy of Sciences. 1966;133(2):391–404. doi: 10.1111/j.1749-6632.1966.tb52379.x. PubMed PMID:5336347.

13. Maczuga N, Tran ENH, Qin J, Morona R. Interdependence of Shigella flexneri O Antigen and Enterobacterial Common Antigen Biosynthetic Pathways. J Bacteriol. 2022;204(4):e0054621. Epub 2022/03/17. doi: 10.1128/jb.00546-21. PubMed PMID: 35293778; PubMed Central PMCID: PMCPMC9017295.

14. Liu D, Reeves PR. Escherichia coli K12 regains its O antigen. Microbiology (Reading). 1994;140 (Pt 1):49–57. Epub 1994/01/01. doi: 10.1099/13500872-140-1-49. PubMed PMID: 7512872.

15. Baba T, Ara T, Hasegawa M, Takai Y, Okumura Y, Baba M, et al. Construction of Escherichia coli K-12 in-frame, single-gene knockout mutants: the Keio collection. Mol Syst Biol. 2006;2:2006 0008. Epub 2006/06/02. doi: 10.1038/msb4100050. PubMed PMID: 16738554; PubMed Central PMCID: PMCPMC1681482.

16. Qin J, Hong Y, Morona R, Totsika M. O antigen biogenesis sensitises Escherichia coli K-12 to bile salts, providing a plausible explanation for its evolutionary loss. PLoS Genet. 2023;19(10):e1010996. Epub 2023/10/04. doi: 10.1371/journal.pgen.1010996. PubMed PMID: 37792901; PubMed Central PMCID: PMCPMC10578602.

17. Datsenko KA, Wanner BL. One-step inactivation of chromosomal genes in Escherichia coli K-12 using PCR products. Proceedings of the National Academy of Sciences of the United States of America. 2000;97(12):6640–5. doi: DOI 10.1073/pnas.120163297. PubMed PMID: WOS:000087526300074.

18. Simon R, Priefer U, Pühler A. A Broad Host Range Mobilization System for In Vivo Genetic Engineering: Transposon Mutagenesis in Gram Negative Bacteria. Bio/Technology. 1983;1(9):784–91. doi: 10.1038/nbt1183-784.

19. Schaller K, Nomura M. Colicin E2 is DNA endonuclease. Proceedings of the National Academy of Sciences of the United States of America. 1976;73(11):3989–93. Epub 1976/11/01. doi: 10.1073/pnas.73.11.3989. PubMed PMID: 1069283; PubMed Central PMCID: PMCPMC431295.

20. Benedetti H, Frenette M, Baty D, Lloubes R, Geli V, Lazdunski C. Comparison of the uptake systems for the entry of various BtuB group colicins into Escherichia coli. J Gen Microbiol. 1989;135(12):3413–20. Epub 1989/12/01. doi: 10.1099/00221287-135-12-3413. PubMed PMID: 2699893.

21. Tran EN, Papadopoulos M, Morona R. Relationship between O-antigen chain length and resistance to colicin E2 in *Shigella flexneri*. Microbiology. 2014;160(Pt 3):589–601. Epub 2014/01/16. doi: 10.1099/mic.0.074955-0. PubMed PMID: 24425769.

22. Yamamoto N, Nakahigashi K, Nakamichi T, Yoshino M, Takai Y, Touda Y, et al. Update on the Keio collection of Escherichia coli single-gene deletion mutants. Mol Syst Biol. 2009;5:335. Epub 2009/12/24. doi: 10.1038/msb.2009.92. PubMed PMID: 20029369; PubMed Central PMCID: PMCPMC2824493.

23. Qin J, Hong Y, Maczuga NT, Morona R, Totsika M. Tolerance mechanisms in polysaccharide biosynthesis: Implications for undecaprenol phosphate recycling in Escherichia coli and Shigella flexneri. PLoS Genet. 2025;21(1):e1011591. Epub 2025/01/30. doi: 10.1371/journal.pgen.1011591. PubMed PMID: 39883743.

24. Perez-Mendoza D, de la Cruz F. Escherichia coli genes affecting recipient ability in plasmid conjugation: are there any? BMC genomics. 2009;10:71. Epub 2009/02/11. doi: 10.1186/1471-2164-10-71. PubMed PMID: 19203375; PubMed Central PMCID: PMCPMC2645431.

25. Szczepaniak J, Webby MN. The Tol Pal system integrates maintenance of the three layered cell envelope. NPJ Antimicrob Resist. 2024;2(1):46. Epub 2025/01/23. doi: 10.1038/s44259-024-00065-0. PubMed PMID: 39843782; PubMed Central PMCID: PMCPMC11721397.

26. Alalam H, Graf FE, Palm M, Abadikhah M, Zackrisson M, Bostrom J, et al. A High-Throughput Method for Screening for Genes Controlling Bacterial Conjugation of Antibiotic Resistance. mSystems. 2020;5(6). Epub 2020/12/29. doi: 10.1128/mSystems.01226-20. PubMed PMID: 33361328; PubMed Central PMCID: PMCPMC7762799.

27. Robledo M, Alvarez B, Cuevas A, Gonzalez S, Ruano-Gallego D, Fernandez LA, et al. Targeted bacterial conjugation mediated by synthetic cell-to-cell adhesions. Nucleic acids research. 2022;50(22):12938–50. Epub 2022/12/14. doi: 10.1093/nar/gkac1164. PubMed PMID: 36511856; PubMed Central PMCID: PMCPMC9825185.

28. Sperandeo P, Pozzi C, Deho G, Polissi A. Non-essential KDO biosynthesis and new essential cell envelope biogenesis genes in the Escherichia coli yrbG-yhbG locus. Res Microbiol. 2006;157(6):547–58. Epub 20060209. doi: 10.1016/j.resmic.2005.11.014. PubMed PMID: 16765569.

29. Chakraborty S, Banerjee P, Joseph JP, Pathak S, Verma T, Karhale AK, et al. Functional loss of rffG and rfbB, encoding dTDP-glucose 4,6-dehydratase, alters colony morphology, cell shape, motility and virulence in Salmonella Typhimurium. Front Microbiol. 2025;16:1572117. Epub 20250521. doi: 10.3389/fmicb.2025.1572117. PubMed PMID: 40469742; PubMed Central PMCID: PMCPMC12136496.

30. Hone DM, Attridge SR, Forrest B, Morona R, Daniels D, LaBrooy JT, et al. A galE via (Vi antigen-negative) mutant of Salmonella typhi Ty2 retains virulence in humans. Infect Immun. 1988;56(5):1326–33. doi: 10.1128/iai.56.5.1326-1333.1988. PubMed PMID: 3356467; PubMed Central PMCID: PMCPMC259821.

31. Mulford CA, Osborn MJ. An intermediate step in translocation of lipopolysaccharide to the outer membrane of Salmonella typhimurium. Proceedings of the National Academy of Sciences of the United States of America. 1983;80(5):1159–63. doi: 10.1073/pnas.80.5.1159. PubMed PMID: 6338498; PubMed Central PMCID: PMCPMC393553.

32. Klein G, Kobylak N, Lindner B, Stupak A, Raina S. Assembly of lipopolysaccharide in Escherichia coli requires the essential LapB heat shock protein. The Journal of biological chemistry. 2014;289(21):14829–53. Epub 20140409. doi: 10.1074/jbc.M113.539494. PubMed PMID: 24722986; PubMed Central PMCID: PMCPMC4031536.

33. Aguiniga LM, Yaggie RE, Schaeffer AJ, Klumpp DJ. Lipopolysaccharide Domains Modulate Urovirulence. Infect Immun. 2016;84(11):3131–40. Epub 20161017. doi: 10.1128/IAI.00315-16. PubMed PMID: 27528276; PubMed Central PMCID: PMCPMC5067746.

34. Feldman MF, Marolda CL, Monteiro MA, Perry MB, Parodi AJ, Valvano MA. The activity of a putative polyisoprenol-linked sugar translocase (Wzx) involved in Escherichia coli O antigen assembly is independent of the chemical structure of the O repeat. J Biol Chem. 1999;274(49):35129–38. doi: 10.1074/jbc.274.49.35129. PubMed PMID: 10574995.

35. Tsai CM, Frasch CE. A sensitive silver stain for detecting lipopolysaccharides in polyacrylamide gels. Anal Biochem. 1982;119(1):115–9. Epub 1982/01/01. doi: 10.1016/0003-2697(82)90673-x. PubMed PMID: 6176137.

36. Sharma O, Datsenko KA, Ess SC, Zhalnina MV, Wanner BL, Cramer WA. Genome-wide screens: novel mechanisms in colicin import and cytotoxicity. Mol Microbiol. 2009;73(4):571–85. Epub 2009/08/05. doi: 10.1111/j.1365-2958.2009.06788.x. PubMed PMID: 19650773; PubMed Central PMCID: PMCPMC3100173.

37. Datsenko KA, Wanner BL. One-step inactivation of chromosomal genes in Escherichia coli K-12 using PCR products. Proceedings of the National Academy of Sciences of the United States of America. 2000;97(12):6640–5. doi: 10.1073/pnas.120163297. PubMed PMID: 10829079; PubMed Central PMCID: PMC18686.

