## Supplementary File for "Genome-wide screen for genes required for smooth lipopolysaccharide production in Escherichia coli K-12"

**Supplementary Table 1.** Low-liquid conjugative mutants found in primary screen.

| No. | Mutant ID [a] | Plate conjugation [b] | Colicin E2 sensitivity [c] |
|---|---|---|---|
| 1 | *sdhD* | +++ | R |
| 2 | *prpC* | +++ | R |
| 3 | *talB* | +++ | R |
| 4 | *atoS* | +++ | R |
| 5 | *phoP* | +++ | R |
| 6 | *cheA* | +++ | R |
| 7 | *rpe* | +++ | S |
| 8 | *pgi* | +++ | R |
| 9 | *pgm* | +++ | R |
| 10 | *rnt* | + (*) | S |
| 11 | *wza* | +++ | R |
| 12 | *dbpA* | +++ | R |
| 13 | *yfaD* | +++ | R |
| 14 | *yeaR* | +++ | R |
| 15 | *ybjC* | +++ | R |
| 16 | *yfbK* | ++ (*) | S |
| 17 | *yedI* | +++ | R |
| 18 | *yfgL* | ++ | R |
| 19 | *uspE* | +++ | R |
| 20 | *yphF* | +++ | R |
| 21 | *yfdH* | + (*) | S |
| 22 | *yagA* | +++ | R |
| 23 | *yciM (lapB)* | +++ | S |
| 24 | *ydcU* | +++ | R |
| 25 | *pncA* | + | S |
| 26 | *yaeJ* | +++ | R |
| 27 | *yafJ* | +++ | S |
| 28 | *yliF* | +++ | R |
| 29 | *gst* | +++ | R |
| 30 | *yqfB* | +++ | R |
| 31 | *yhcA* | +++ | R |
| 32 | *yjhP* | +++ | R |
| 33 | *brnQ* | +++ | R |
| 34 | *sixA* | +++ | R |
| 35 | *moaD* | +++ | S |
| 36 | *wzxB (rfbX)* | +++ | R |
| 37 | *rfe* | ++ (*) | S |
| 38 | *ycjU* | + (*) | S |

| | | | |
|---|---|---|---|
| 39 | *gmhA (lpcA)* | +++ | S |
| 40 | *xseA* | +++ | R |
| 41 | *glnH* | +++ | R |
| 42 | *fliR* | +++ | R |
| 43 | *tolR* | +++ | R |
| 44 | *torD* | +++ | R |
| 45 | *tolB* | +++ | R |
| 46 | *eptA* | +++ | R |
| 47 | *yjfF* | ++ | S |
| 48 | *slyA* | + | S |
| 49 | *yfcE* | + | S |
| 50 | *ygeN* | ++ (*) | S |
| 51 | *yegK* | +++ | R |
| 52 | *yehD* | +++ | R |
| 53 | *mhpB* | +++ | R |
| 54 | *phnO* | ++ | R |
| 55 | *rbfA* | +++ | R |
| 56 | *csgC* | +++ | R |
| 57 | *dnaT* | +++ | S |
| 58 | *yjcQ* | +++ | R |
| 59 | *afuB* | +++ | R |
| 60 | *yahA* | +++ | R |
| 61 | *yajC* | +++ | R |
| 62 | *essD* | +++ | R |
| 63 | *ybeD* | +++ | R |
| 64 | *ybgK* | +++ | R |
| 65 | *pal* | +++ | R |
| 66 | *ihfA* | +++ | R |
| 67 | *rpmI* | +++ | R |
| 68 | *ruvC* | +++ | R |
| 69 | *yeiQ* | +++ | R |
| 70 | *ompC* | +++ | R |
| 71 | *frlA* | +++ | R |
| 72 | *waaG (rfaG)* | +++ | S |
| 73 | *waaQ (rfaQ)* | +++ | R |
| 74 | *rffA* | +++ | R |
| 75 | *cpxA* | +++ | R |
| 76 | *trmC* | +++ | R |
| 77 | *rimM* | +++ | R |
| 78 | *ypjJ* | +++ | R |
| 79 | *xdhA* | +++ | R |
| 80 | *ygfB* | +++ | R |
| 81 | *dcrB* | +++ | R |

| | | | |
|---|---|---|---|
| 82 | *rnr* | +++ | R |
| 83 | *dedD* | +++ | R |
| 84 | *ompX* | +++ | R |
| 85 | *yhjG* | +++ | R |

[a] Mutants highlighted in green indicate additional CSMs discovered after plate conjugations; Mutants highlighted in yellow indicate low-conjugative mutants.
[b] Conjugation scores based on growth of conjugate dot on plates as determined by eye; +++ (similar to control BW25113 [pSUP203-WbbL] strain), ++/+ (less than control), - (no growth detected). Mutants requiring conjugate mix to be concentrated prior to dotting for observable growth on plates are starred in brackets.
[c] Conjugate sensitivity on plates containing 0.2 µg/ml colicin E2; R (resistant), S (sensitive).

**Supplementary Table 2. List of colicin sensitive mutants.**

| No. | Mutant ID | Protein function in *E. coli* | LPS profile (+ pSUP203-WbbL) |
|---|---|---|---|
| 1 | *cytR* | DNA-binding transcriptional repressor CytR | S-LPS |
| 2 | *sdhC* | Succinate:quinone oxidoreductase, membrane protein SdhC | Reduced Oag/Lipid A-Core defect |
| 3 | *deoR* | DeoR DNA-binding transcriptional repressor DeoR | S-LPS |
| 4 | *argP* | DNA-binding transcriptional dual regulator ArgP | R-LPS |
| 5 | *nanR* | DNA-binding transcriptional dual regulator NanR | S-LPS |
| 6 | *sdhA* | Succinate:quinone oxidoreductase, FAD binding protein | S-LPS |
| 7 | *cyoD* | Cytochrome *bo*$_3$ ubiquinol oxidase subunit 4 | S-LPS |
| 8 | *envY* | DNA-binding transcriptional activator EnvY | S-LPS |
| 9 | *sdiA* | DNA-binding transcriptional dual regulator SdiA | S-LPS |
| 10 | *csgD* | DNA-binding transcriptional dual regulator CsgD | S-LPS |
| 11 | *fadR* | DNA-binding transcriptional dual regulator FadR | S-LPS |
| 12 | *nac* | DNA-binding transcriptional dual regulator Nac | S-LPS |
| 13 | *hupA* | Transcriptional dual regulator HU-α (HU-2) | S-LPS |
| 14 | *malI* | DNA-binding transcriptional repressor MalI | S-LPS |
| 15 | *ybeF* | Putative LysR-type DNA-binding transcriptional regulator YbeF | S-LPS |
| 16 | *fdrA* | Putative acyl-CoA synthetase FdrA | S-LPS |
| 17 | *gnsA* | predicted regulator of phosphatidylethanolamine synthesis | S-LPS |

| | | | |
|---|---|---|---|
| 18 | *ycbU* | predicted fimbrial-like adhesin protein | Reduced Oag/Core defect |
| 19 | *rfaE (hldE)* | fused heptose 7-phosphate kinase/heptose 1-phosphate adenyltransferase | No Oag/Lipid A-Core defect |
| 20 | *ydcS* | putative transport protein, ABC superfamily - periplasmic binding protein | S-LPS |
| 21 | *wzc* | protein-tyrosine kinase | Reduced Oag/Lipid A-Core defect |
| 22 | *hfq* | HF-I, host factor for RNA phage Q β replication | S-LPS |
| 23 | *adhE* | Fused acetaldehyde-CoA dehydrogenase and iron-dependent alcohol dehydrogenase | S-LPS |
| 24 | *mdh* | Malate dehydrogenase | S-LPS |
| 25 | *cheB* | Protein-glutamate methylesterase/protein glutamine deamidase | S-LPS |
| 26 | *nuoA* | NADH:quinone oxidoreductase subunit A | S-LPS |
| 27 | *citA (dpiB)* | Sensor histidine kinase DpiB | S-LPS |
| 28 | *uvrY* | DNA-binding transcriptional activator UvrY | S-LPS |
| 29 | *evgA* | DNA-binding transcriptional activator EvgA | S-LPS |
| 30 | *gapC* | Glyceraldehyde-3-phosphate dehydrogenase | S-LPS |
| 31 | *narX* | Sensor histidine kinase NarX | S-LPS |
| 32 | *yedV (hprS)* | Sensor histidine kinase HprS | Reduced Oag |
| 33 | *evgS* | Sensor histidine kinase EvgS | S-LPS |
| 34 | *fumA* | Fumarase A | S-LPS |
| 35 | *phoB* | DNA-binding transcriptional dual regulator PhoB | S-LPS |
| 36 | *cusR* | DNA-binding transcriptional activator CusR | S-LPS |
| 37 | *ldhA* | D-lactate dehydrogenase | Reduced Oag |
| 38 | *gltA* | Citrate synthase | S-LPS |
| 39 | *yfdV* | Putative transport protein YfdV | S-LPS |
| 40 | *lpd* | Lipoamide dehydrogenase | S-LPS |
| 41 | *ynfL* | Putative DNA-binding transcriptional regulator | S-LPS |
| 42 | *mrcB* | Penicillin binding protein 1B (PBP1B) | S-LPS |
| 43 | *ompA* | Outer membrane protein A | S-LPS |
| 44 | *lhr* | ATP-dependent helicase Lhr | S-LPS |
| 45 | *potH* | Putrescine ABC transporter membrane subunit PotH | S-LPS |
| 46 | *lon* | Lon protease | S-LPS |
| 47 | *ydcP (rlhA)* | 23S rRNA 5-hydroxycytidine C2501 synthase | S-LPS |
| 48 | *srlR* | GutR DNA-binding transcriptional repressor | S-LPS |
| 49 | *acnA* | aconitate hydratase 1 | S-LPS |

| | | | |
|---|---|---|---|
| 50 | *frdC* | fumarate reductase membrane protein | S-LPS |
| 51 | *pykA* | pyruvate kinase II | S-LPS |
| 52 | *aceA* | isocitrate lyase | S-LPS |
| 53 | *appC* | cytochrome bd-II terminal oxidase subunit I | S-LPS |
| 54 | *dps* | Dps complex; stationary phase nucleoid component that sequesters iron & protects DNA from damage | S-LPS |
| 55 | *phoR* | PhoR sensory histidine kinase | S-LPS |
| 56 | *torR* | TorR transcriptional dual regulator | S-LPS |
| 57 | *poxB* | pyruvate oxidase | S-LPS |
| 58 | *tktA* | transketolase I | S-LPS |
| 59 | *rbsB* | ribose ABC transporter - putative periplasmic binding protein | S-LPS |
| 60 | *hchA* | glyoxalase III, Hsp31 molecular chaperone | S-LPS |
| 61 | *yfdF* | predicted protein | S-LPS |
| 62 | *glgC* | glucose-1-phosphate adenylyltransferase | S-LPS |
| 63 | *ppk* | polyphosphate kinase | S-LPS |
| 64 | *arcB* | ArcB sensory histidine kinase | S-LPS |
| 65 | *ackA* | acetate kinase | S-LPS |
| 66 | *hofQ* | protein involved in utilization of DNA as a carbon source | S-LPS |
| 67 | *basS* | BasS sensory histidine kinase | S-LPS |
| 68 | *ybhD* | predicted DNA-binding transcriptional regulator | S-LPS |
| 69 | *tesA* | multifunctional acyl-CoA thioesterase I and protease I and lysophospholipase L1 | S-LPS |
| 70 | *pepT* | peptidase T | S-LPS |
| 71 | *degP* | serine protease Do | S-LPS |
| 72 | *dacA* | D-alanyl-D-alanine carboxypeptidase, fraction A; penicillin-binding protein 5 | S-LPS |
| 73 | *yeaY* | predicted lipoprotein | S-LPS |
| 74 | *yeiE* | LYSR-type transcriptional regulator | S-LPS |
| 75 | *yqeI* | predicted transcriptional regulator | S-LPS |
| 76 | *srmB* | SrmB, DEAD-box RNA helicase | S-LPS |
| 77 | *ygeF* | predicted protein | S-LPS |
| 78 | *yfbP* | predicted protein | S-LPS |
| 79 | *mepA* | murein DD-endopeptidase, penicillin-insensitive | S-LPS |
| 80 | *yfaP* | conserved protein | S-LPS |
| 81 | *yfbR* | dCMP phosphohydrolase | S-LPS |
| 82 | *htpX* | heat shock protein, integral membrane protein | S-LPS |
| 83 | *yhbU* | predicted peptidase (collagenase-like) | S-LPS |
| 84 | *yfaQ* | predicted protein | S-LPS |
| 85 | *yfeZ* | predicted inner membrane protein | S-LPS |
| 86 | *degQ* | serine endoprotease, periplasmic | S-LPS |

| | | | |
|---|---|---|---|
| 87 | *yfbH (arnD)* | putative 4-deoxy-4-formamido-L-arabinose-phosphoundecaprenol deformylase ArnD | S-LPS |
| 88 | *yfdX* | predicted protein | S-LPS |
| 89 | *hycH* | protein required for maturation of hydrogenase 3 | S-LPS |
| 90 | *ypeC* | conserved protein | S-LPS |
| 91 | *eutT* | predicted cobalamine adenosyltransferase in ethanolamine utilization | S-LPS |
| 92 | *yegQ* | tRNA wobble base hydroxylation protein TrhP | R-LPS |
| 93 | *hybD* | predicted maturation peptidase for hydrogenase 2 | S-LPS |
| 94 | *hslU* | HslU hexamer | S-LPS |
| 95 | *yfjK (abpB)* | CP4-57 prophage; anti-bacteriophage protein; mutations lead to radiation resistance | S-LPS |
| 96 | *ydhT* | conserved protein | S-LPS |
| 97 | *yoaH* | conserved protein | S-LPS |
| 98 | *ypfE (eutS)* | predicted structural protein, ethanolamine utilization microcompartment | S-LPS |
| 99 | *ypjF* | CP4-57 prophage; toxin of the YpjF-YfjZ toxin-antitoxin system | S-LPS |
| 100 | *yfjN (rnlA)* | CP4-57 prophage; RNase LS, toxin of the RnlAB toxin-antitoxin system | S-LPS |
| 101 | *yeaC* | conserved protein | S-LPS |
| 102 | *yecE* | conserved protein | S-LPS |
| 103 | *ynfB* | predicted protein | S-LPS |
| 104 | *ydfU* | Qin prophage; predicted protein | S-LPS |
| 105 | *chbG* | chito-oligosaccharide mono-deacetylase | S-LPS |
| 106 | *yobB* | conserved protein | S-LPS |
| 107 | *yecR* | predicted protein | S-LPS |
| 108 | *ycfT* | inner membrane protein YcfT | S-LPS |
| 109 | *ycjY* | predicted hydrolase | S-LPS |
| 110 | *ydcA* | predicted protein | S-LPS |
| 111 | *ydbH* | putative intermembrane phospholipid transport protein | S-LPS |
| 112 | *ydcE (pptA)* | probable 4-oxalocrotonate tautomerase (4-OT) | S-LPS |
| 113 | *yahG* | conserved protein | S-LPS |
| 114 | *yejK* | nucleoid-associated protein | S-LPS |
| 115 | *yfaX* | predicted DNA-binding transcriptional regulator | S-LPS |
| 116 | *ybgL* | predicted lactam utilization protein | S-LPS |
| 117 | *ycbC (elyC)* | inner membrane protein involved in peptidoglycan synthesis | S-LPS |
| 118 | *ynfA* | conserved inner membrane protein | S-LPS |
| 119 | *yeaL* | conserved inner membrane protein | S-LPS |
| 120 | *yeeS* | CP4-44 prophage; predicted DNA repair protein | S-LPS |

| | | | |
|---|---|---|---|
| 121 | *yedK* | predicted protein | S-LPS |
| 122 | *yedA* | putative transport protein, drug/metabolite exporter family | S-LPS |
| 123 | *ybbY* | putative transport protein, nucleobase:cation symporter-2 (NCS2) family | S-LPS |
| 124 | *ybhB* | predicted kinase inhibitor | S-LPS |
| 125 | *ycbK* | conserved protein | S-LPS |
| 126 | *yfaO (nudI)* | pyrimidine deoxynucleoside triphosphate pyrophosphohydrolase | S-LPS |
| 127 | *ycjG* | L-Ala-D/L-Glu epimerase | S-LPS |
| 128 | *ybgC* | esterase/thioesterase | S-LPS |
| 129 | *yccM* | predicted 4Fe-4S membrane protein | S-LPS |
| 130 | *ychP (ychO)* | predicted invasin | S-LPS |
| 131 | *yqcA* | predicted flavoprotein | S-LPS |
| 132 | *lomR* | Rac prophage; predicted protein, N-ter fragment | S-LPS |
| 133 | *yafE* | predicted S-adenosylmethionine-dependent methyltransferase | S-LPS |
| 134 | *ybjJ* | Inner membrane protein YbjJ | Reduced Oag |
| 135 | *mprA* | DNA-binding transcriptional repressor MprA | No Oag/Core-Lipid A defect |
| 136 | *cof* | pyridoxal phosphatase / HMP-PP hydrolase | S-LPS |
| 137 | *yfcY (fadI)* | Fad I component of anaerobic fatty acid oxidation complex | S-LPS |
| 138 | *flk* | predicted flagella assembly protein | S-LPS |
| 139 | *ypdE* | broad-specificity exoaminopeptidase | S-LPS |
| 140 | *wcaK* | predicted colanic acid biosynthesis pyruvyl transferase | S-LPS |
| 141 | *ycbS (elfC)* | predicted outer membrane usher protein | S-LPS |
| 142 | *yiiQ* | conserved protein | S-LPS |
| 143 | *ydjJ* | predicted oxidoreductase, Zn-dependent and NAD(P)-binding | S-LPS |
| 144 | *ydfH* | YdfH DNA-binding transcriptional repressor | S-LPS |
| 145 | *ymcA (gfcD)* | putative lipoprotein | S-LPS |
| 146 | *pepP* | proline aminopeptidase P II | S-LPS |
| 147 | *ydjZ* | conserved inner membrane protein | S-LPS |
| 148 | *ydiJ* | predicted FAD-linked oxidoreductase | S-LPS |
| 149 | *sufC* | SufC component of SufBCD Fe-S cluster scaffold complex | S-LPS |
| 150 | *ygfZ* | folate-binding protein | S-LPS |
| 151 | *yghZ* | L-glyceraldehyde 3-phosphate reductase | S-LPS |

| | | | |
|---|---|---|---|
| 152 | *yqjA* | membrane transport protein; required for proton-motive force (PMF) dependent drug efflux | S-LPS |
| 153 | *ybdR* | predicted oxidoreductase, Zn-dependent and NAD(P)-binding | S-LPS |
| 154 | *htrG (ygiM)* | predicted signal transduction protein (SH3 domain) | S-LPS |
| 155 | *yhcO* | predicted barnase inhibitor | S-LPS |
| 156 | *hdeA* | HdeA dimer, inactive form of acid-resistance protein | S-LPS |
| 157 | *ysaB* | predicted protein | S-LPS |
| 158 | *damX* | cell division protein DamX | S-LPS |
| 159 | *yrbC (mlaC)* | phospholipid ABC transporter - periplasmic binding protein | S-LPS |
| 160 | *bax* | conserved protein | S-LPS |
| 161 | *yhdH* | acrylyl-CoA reductase | S-LPS |
| 162 | *yhfG* | DUF2559 domain-containing protein YhfG | Reduced Oag |
| 163 | *yhiN* | predicted oxidoreductase with FAD/NAD(P)-binding domain | S-LPS |
| 164 | *yhdT* | conserved inner membrane protein | S-LPS |
| 165 | *yhgE* | putative transport protein, YhgE family | S-LPS |
| 166 | *yhjB* | predicted DNA-binding response regulator in two-component regulatory system | S-LPS |
| 167 | *yicC* | conserved protein | S-LPS |
| 168 | *yjaG* | conserved protein | S-LPS |
| 169 | *yjeB (nsrR)* | NsrR DNA-binding transcriptional repressor | S-LPS |
| 170 | *rffC (wecD)* | dTDP-4-amino-4,6-dideoxy-D-galactose acyltransferase | Reduced Oag |
| 171 | *yihU* | 3-sulfolactaldehyde reductase | S-LPS |
| 172 | *yiiX* | conserved hypothetical protein of the NlpC/P60 peptidase superfamily | S-LPS |
| 173 | *yigA* | conserved protein | S-LPS |
| 174 | *hofC* | inner membrane protein HofC | S-LPS |
| 175 | *ulaR* | UlaR DNA-binding transcriptional repressor | S-LPS |
| 176 | *uvrA* | excision nuclease subunit A | S-LPS |
| 177 | *ydaV* | Rac prophage; predicted DNA replication protein | S-LPS |
| 178 | *holC* | DNA polymerase III, $\chi$ subunit | S-LPS |
| 179 | *lysC* | aspartate kinase III | S-LPS |
| 180 | *aroL* | shikimate kinase II | S-LPS |
| 181 | *lipA* | lipoyl synthase | S-LPS |
| 182 | *moaE* | molybdopterin synthase large subunit | S-LPS |
| 183 | *rfaP (waaP)* | Lipopolysaccharide core heptose (I) kinase | Reduced Oag/Core defect |

| | | | |
|---|---|---|---|
| 184 | *rfaD* | ADP-L-glycero-D-mannoheptose 6-epimerase | No Oag/Core defect |
| 185 | *rffH* | dTDP-glucose pyrophosphorylase (involved in ECA biosynthesis) | S-LPS |
| 186 | *motA* | MotA protein, proton conductor component of motor; no effect on switching | S-LPS |
| 187 | *rfaF (waaF)* | ADP-heptose—LPS heptosyltransferase 2 | No Oag/Core defect |
| 188 | *rfaL (waaL)* | O-antigen ligase | No Oag/Core defect |
| 189 | *dnaK* | chaperone protein DnaK | S-LPS |
| 190 | *ccmG* | holocytochrome c synthetase - thiol:disulfide oxidoreductase CcmG | S-LPS |
| 191 | *dnaJ* | chaperone protein DnaJ | S-LPS |
| 192 | *lpxM* | Lipid A biosynthesis myristoyltransferase | No Oag/Core defect |
| 193 | *rfaJ (waaJ)* | UDP-glucose:(glucosyl)LPS α-1,2-glucosyltransferase | No Oag/Core defect |
| 194 | *oppF* | Murein tripeptide ABC transporter / oligopeptide ABC transporter ATP binding subunit OppF | No Oag |
| 195 | *rfaI (waaO)* | UDP-D-glucose:(glucosyl)LPS α-1,3-glucosyltransferase | No Oag/Lipid A-Core defect |
| 196 | *proX* | glycine betaine / proline ABC transporter - periplasmic binding protein | S-LPS |
| 197 | *rfbD (rmlD)* | dTDP-4-dehydrorhamnose reductase | No Oag/Core defect |
| 198 | *rfbB* | dTDP-glucose 4,6-dehydratase 1 | S-LPS |
| 199 | *nagA* | N-acetylglucosamine-6-phosphate deacetylase | S-LPS |
| 200 | *cpsG* | Phosphomannomutase | S-LPS |
| 201 | *rfbC* | dTDP-4-dehydrorhamnose 3,5-epimerase | No Oag/Core defect |
| 202 | *rfbA* | Glucose-1-phosphate thymidylyltransferase 1 | Reduced Oag |
| 203 | *nfnB (nfsB)* | NAD(P)H nitroreductase NfsB | S-LPS |
| 204 | *galU* | UTP—glucose-1-phosphate uridylyltransferase | No Oag/Core defect |
| 205 | *galE* | UDP-glucose 4-epimerase | Reduced Oag |
| 206 | *yiaI (ysaA)* | predicted hydrogenase, 4Fe-4S ferredoxin-type component | S-LPS |
| 207 | *yhaJ* | predicted DNA-binding transcriptional regulator LYSR-type | S-LPS |
| 208 | *phnB (yjdN)* | conserved protein | S-LPS |
| 209 | *dcd* | dCTP deaminase | S-LPS |
| 210 | *araE* | arabinose:H+ symporter | S-LPS |
| 211 | *cyaA* | adenylate cyclase | S-LPS |

| | | | |
|---|---|---|---|
| 212 | *ddpD* | YddP: an ATP-binding component of a predicted peptide ABC transporter | S-LPS |
| 213 | *malX* | MalX PTS permease | S-LPS |
| 214 | *glnQ* | glutamine ABC transporter - ATP binding subunit | S-LPS |
| 215 | *btuE* | thioredoxin/glutathione peroxidase | S-LPS |
| 216 | *lysP* | lysine:H+ symporter | S-LPS |
| 217 | *queA* | S-adenosylmethionine:tRNA ribosyltransferase-isomerase | S-LPS |
| 218 | *exbB* | TonB energy transducing system - ExbB subunit (ivolved in transport of iron & Vit. B12 across OM) | S-LPS |
| 219 | *ytfF* | inner membrane protein YtfF | S-LPS |
| 220 | *kbaY* | KbaY (no known function) | S-LPS |
| 221 | *uraA* | uracil:H+ symporter UraA | S-LPS |
| 222 | *gudP* | galactarate / glucarate / glycerate transporter GudP | S-LPS |
| 223 | *pitB* | metal phosphate:H+ symporter PitB | S-LPS |
| 224 | *elaC (rbn)* | RNase BN | S-LPS |
| 225 | *sbcB* | exonuclease I, 3' --> 5' specific; deoxyribophosphodiesterase | S-LPS |
| 226 | *xylH* | xylose ABC transporter - membrane subunit | S-LPS |
| 227 | *ptsN* | phosphotransferase system enzyme IIA, regulation of potassium transport | S-LPS |
| 228 | *yfaV* | putative transport protein, major facilitator superfamily | S-LPS |
| 229 | *dcuD* | putative transport protein, C4-dicarboxylate uptake C family | S-LPS |
| 230 | *puuP* | putrescine:H+ symporter PuuP | S-LPS |
| 231 | *hyfA* | hydrogenase 4, component A | S-LPS |
| 232 | *folP* | dihydropteroate synthase | S-LPS |
| 233 | *dtd* | D-Tyr-tRNATyr deacylase | S-LPS |
| 234 | *rpsF* | 30S ribosomal subunit protein S6 | S-LPS |
| 235 | *rpmJ* | 50S ribosomal subunit protein L36 | S-LPS |
| 236 | *frlA* | fructoselysine / psicoselysine transporter | S-LPS |
| 237 | *alaS* | alanine—tRNA ligase/DNA-binding transcriptional repressor | S-LPS |
| 238 | *selB* | selenocysteyl-tRNA-specific translation elongation factor | S-LPS |
| 239 | *cspG* | cold shock protein CspG | S-LPS |
| 240 | *yiiT (uspD)* | stress protein involved in resistance to UV irradiation | S-LPS |
| 241 | *surE (umpG)* | broad specificity 5'(3')-nucleotidase and polyphosphatase | S-LPS |

| | | | |
|---|---|---|---|
| 242 | *pspE* | thiosulfate sulfurtransferase | S-LPS |
| 243 | *fecD* | ferric dicitrate ABC transporter - membrane subunit | S-LPS |
| 244 | *ligB* | DNA ligase | S-LPS |
| 245 | *phnE* | phosphonate ABC transporter - membrane subunit | S-LPS |
| 246 | *yjjQ* | YjjQ DNA-binding transcriptional repressor | S-LPS |
| 247 | *dsbA* | protein disulfide oxidoreductase - DsbA(reduced) | S-LPS |
| 248 | *yibD (waaH)* | UDP-glucuronate:LPS(HepIII) glycosyltransferase | Reduced Oag |
| 249 | *ppiC* | peptidyl-prolyl cis-trans isomerase C (rotamase C) | S-LPS |
| 250 | *yihP* | putative 2,3-dihydroxypropane-1-sulfonate export protein | S-LPS |
| 251 | *nmpC* | outer membrane porin protein; locus of qsr prophage | S-LPS |
| 252 | *alx* | predicted membrane-bound redox modulator that is induced by high pH | S-LPS |
| 253 | *ptsA* | fused predicted PTS enzyme : HPr component / enzyme I component / enzyme IIA component | S-LPS |
| 254 | *yjbO (pspG)* | phage shock protein G | S-LPS |
| 255 | *yjjA* | conserved protein | S-LPS |
| 256 | *ycdL (rutB)* | peroxyureidoacrylate / ureidoacrylate amido hydrolase | S-LPS |
| 257 | *ybhJ* | predicted hydratase | S-LPS |
| 258 | *yaiU* | predicted protein | S-LPS |
| 259 | *ycgN* | conserved protein | Reduced Oag |
| 260 | *yecN* | predicted inner membrane protein | S-LPS |
| 261 | *yegX* | predicted hydrolase | S-LPS |
| 262 | *tolC* | TolC outer membrane channel | Reduced Oag |
| 263 | *modA* | molybdate ABC transporter - periplasmic binding protein | S-LPS |
| 264 | *glpR* | GlpR DNA-binding transcriptional repressor | S-LPS |
| 265 | *xerC* | site-specific tyrosine recombinase | S-LPS |
| 266 | *gmhB* | D,D-heptose 1,7-bisphosphate phosphatase (involved in LPS core synthesis) | Reduced Oag/Core defect |
| 267 | *rsxC* | member of SoxR-reducing complex | S-LPS |
| 268 | *dam* | DNA adenine methyltransferase | Reduced Oag |
| 269 | *yfdS* | CPS-53 (KpLE1) prophage; predicted protein | S-LPS |
| 270 | *rfaC (waaC)* | ADP-heptose:LPS heptosyltransferase I | No Oag/Core defect |
| 271 | *bglH* | carbohydrate-specific outer membrane porin, cryptic | S-LPS |
| 272 | *guaA* | GMP synthetase | S-LPS |

| | | | |
|---|---|---|---|
| 273 | *recD* | polypeptide RecD | S-LPS |
| 274 | *pcnB* | poly(A) polymerase I | S-LPS |
| 275 | *yeeY* | predicted DNA-binding transcriptional regulator | S-LPS |
| 276 | *cld (wzzB)* | regulator of length of O-antigen component of lipopolysaccharide chains | Unregulated Oag |
| 277 | *yiiM* | protein involved in base analog detoxification | S-LPS |
| 278 | *nuoC* | NADH:ubiquinone oxidoreductase, chain CD | S-LPS |
| 279 | *ynfC* | YnfC lipoprotein | S-LPS |
| 280 | *yeeP* | CP4-44 prophage; predicted GTP-binding protein | S-LPS |
| 281 | *tolA* | TolA - inner membrane protein of the Tol-Pal system | S-LPS |
| 282 | *ybiW* | predicted pyruvate formate lyase | S-LPS |
| 283 | *aaeA* | hydroxylated, aromatic carboxylic acid efflux transporter - putative membrane fusion protein | S-LPS |
| 284 | *crp* | CRP transcriptional dual regulator | Reduced Oag |
| 285 | *rfaH* | RfaH transcriptional antiterminator | No Oag/Core defect |
| 286 | *tpiA* | triosephosphate isomerase | S-LPS |
| 287 | *sucA* | 2-oxoglutarate decarboxylase, thiamine-requiring | S-LPS |
| 288 | *phoQ* | PhoQ sensory histidine kinase | S-LPS |
| 289 | *ybeA (rlmH)* | 23S rRNA m3Ψ1915 methyltransferase | S-LPS |
| 290 | *pitA* | metal phosphate:H+ symporter PitA | S-LPS |
| 291 | *aroD* | 3-dehydroquinate dehydratase | S-LPS |
| 292 | *ycdN (efeU)* | hypothetical protein of the OFeT transport family | S-LPS |
| 293 | *ypjL (ypjM_3)* | CP4-57 prophage; predicted inner membrane protein | S-LPS |
| 294 | *eutJ* | predicted chaperonin, ethanolamine utilization protein | S-LPS |
| 295 | *yfjU* | CP4-57 prophage; conserved protein | S-LPS |
| 296 | *yhdV* | predicted outer membrane lipoprotein | S-LPS |
| 297 | *yiiG* | conserved protein | S-LPS |
| 298 | *sfmF* | predicted fimbrial-like adhesin protein | S-LPS |
| 299 | *yghS* | predicted protein with nucleoside triphosphate hydrolase domain | S-LPS |
| 300 | *yfdI* | CPS-53 (KpLE1) prophage; predicted inner membrane protein | Shifted Oag modality |
| 301 | *ynjB* | conserved protein | S-LPS |
| 302 | *wbbK* | predicted lipopolysaccharide biosynthesis protein | No Oag/Core defect |
| 303 | *wbbI* | β-1,6-galactofuranosyltransferase | No Oag/Core defect |
| 304 | *rfaB (waaB)* | UDP-D-galactose:(glucosyl)lipopolysaccharide-1,6-D-galactosyltransferase | Reduced Oag/Core defect |
| 305 | *wbbJ* | predicted acyl transferase | No Oag/Core defect |

| | | | |
|---|---|---|---|
| 306 | *glf* | UDP-galactopyranose mutase (required for Oag biosynthesis) | No Oag/Core defect |
| 307 | *yhhI* | predicted transposase | S-LPS |
| 308 | *nhaA* | Na+:H+ antiporter NhaA | S-LPS |
| 309 | *rpe* | ribulose-phosphate 3-epimerase | S-LPS |
| 310 | *yciM (lapB)* | lipopolysaccharide assembly protein B | No Oag/Lipid A-Core defect |
| 311 | *pncA* | nicotinamidase | S-LPS |
| 312 | *yafJ* | putative glutamine amidotransferase YafJ | S-LPS |
| 313 | *moaD* | molybdopterin synthase sulfur carrier subunit | S-LPS |
| 314 | *lpcA (gmhA)* | D-sedoheptulose 7-phosphate isomerase | No Oag/Lipid A-Core defect |
| 315 | *yjfF* | galactofuranose ABC transporter putative membrane subunit YjtF | S-LPS |
| 316 | *slyA* | DNA-binding transcriptional dual regulator SlyA | Reduced Oag |
| 317 | *yfcE* | phosphodiesterase YfcE | No Oag/Lipid A-Core defect |
| 318 | *dnaT* | primosomal protein DnaT | S-LPS |
| 319 | *rfaG (waaG)* | lipopolysaccharide glucosyltransferase I | DR-LPS |

[a] Mutants highlighted in orange are known to be associated with LPS biosynthesis.

[d] Mutants highlighted in yellow indicate CSMs displaying a defect in S-LPS profile when compared to the parent BW25113 [pSUP203-WbbL] strain, with no known association to S-LPS production.

**Table S3.** Validation of S-LPS defective KEIO mutants (18 mutants)

| **Mutant ID** | **Protein function in *E. coli*** | **LPS** [+ pSUP203-WbbL] | **Re-made mutant LPS** [+ pSUP203-WbbL] | **WGS** |
|---|---|---|---|---|
| *sdhC* | Succinate:quinone oxidoreductase, membrane protein SdhC | DR-LPS | nd | *rcsC::kan* mutant with secondary *gntX* mutation, insertion in *waaB* |
| *argP* | DNA-binding transcriptional dual regulator ArgP | R-LPS | S-LPS | *argP::kan* mutant with secondary *lrhA, wbbJ* mutations |
| *yedV (hprS)* | Sensor histidine kinase HprS | Reduced Oag | S-LPS | nd |
| *ldhA* | D-lactate dehydrogenase | Reduced Oag | S-LPS | nd |

| | | | | |
|---|---|---|---|---|
| *yegQ* | tRNA wobble base hydroxylation protein TrhP | R-LPS | S-LPS | *yegQ::kan* mutant, insertion in *glf* |
| *ybjJ* | Inner membrane protein YbjJ | Reduced Oag | S-LPS | *ybjJ::kan* mutant with secondary *aceE* mutation |
| *mprA* | DNA-binding transcriptional repressor MprA | DR-LPS | S-LPS | *mprA::kan* mutant with secondary *rseB*, *yraH* mutations, insertion in *waaC* |
| *yhfG* | DUF2559 domain-containing protein YhfG | Reduced Oag | S-LPS | nd |
| *lpxM* | Lipid A biosynthesis myristoyltransferase | DR-LPS | S-LPS | nd |
| *oppF* | Murein tripeptide ABC transporter / oligopeptide ABC transporter ATP binding subunit OppF | SR-LPS | nd | No *kan* insertion, secondary *wzyB* mutation |
| *ycgN* | Putative metal-chelating domain-containing protein YcgN | Reduced Oag | S-LPS | nd |
| *tolC* | TolC outer membrane channel | Reduced Oag | S-LPS | nd |
| *dam* | DNA adenine methyltransferase | Reduced Oag | S-LPS | nd |
| *crp* | CRP transcriptional dual regulator | Reduced Oag | S-LPS | nd |
| *wzc* | Protein-tyrosine kinase | DR-LPS* (with reduced Oag) | S-LPS | *wzc::kan* mutant with secondary *gmhB* mutation |
| *ycbU* | Predicted fimbrial-like adhesin protein | DR-LPS* (with reduced Oag) | S-LPS | *ycbU::kan* with secondary *selB*, *aceE* mutations |
| *slyA* | DNA-binding transcriptional dual regulator SlyA | Reduced Oag | S-LPS | nd |
| *yfcE* | Phosphodiesterase YfcE | DR-LPS | S-LPS | nd |
